# Colonization and Dissemination of *Klebsiella pneumoniae* is Dependent on Dietary Carbohydrates

**DOI:** 10.1101/2023.05.25.542283

**Authors:** Aaron L. Hecht, Lisa C. Harling, Elliot S. Friedman, Ceylan Tanes, Junhee Lee, Jenni Firrman, Vincent Tu, LinShu Liu, Kyle Bittinger, Mark Goulian, Gary D. Wu

**Affiliations:** Division of Gastroenterology and Hepatology, Hospital of the University of Pennsylvania, Philadelphia, PA; Division of Gastroenterology, Hepatology, and Nutrition, The Children’s Hospital of Philadelphia, Philadelphia, PA; Department of Biology, University of Pennsylvania, Philadelphia, PA; Dairy and Functional Foods Research Unit, Eastern Regional Research Center, Agricultural Research Service, US Department of Agriculture, Wyndmoor, PA

## Abstract

Dysbiosis of the gut microbiota is increasingly appreciated as both a consequence and precipitant of human disease. The outgrowth of the bacterial family *Enterobacteriaceae* is a common feature of dysbiosis, including the human pathogen *Klebsiella pneumoniae*. Dietary interventions have proven efficacious in the resolution of dysbiosis, though the specific dietary components involved remain poorly defined. Based on a previous human diet study, we hypothesized that dietary nutrients serve as a key resource for the growth of bacteria found in dysbiosis. Through human sample testing, and *ex-vivo*, and *in vivo* modeling, we find that nitrogen is not a limiting resource for the growth of *Enterobacteriaceae* in the gut, contrary to previous studies. Instead, we identify dietary simple carbohydrates as critical in colonization of *K. pneumoniae*. We additionally find that dietary fiber is necessary for colonization resistance against *K. pneumoniae*, mediated by recovery of the commensal microbiota, and protecting the host against dissemination from the gut microbiota during colitis. Targeted dietary therapies based on these findings may offer a therapeutic strategy in susceptible patients with dysbiosis.

## Introduction

The gut microbiome is increasingly appreciated to play a key role in human disease. Dysbiosis, an alteration of the microbiome associated with chronic disease, is often found in patients with inflammatory bowel disease (IBD) and liver cirrhosis^1–4^. Common features of dysbiosis that characterize IBD and cirrhosis are reduced bacterial diversity and the outgrowth of *Enterobacteriaceae*, including *Escherichia coli* and *Klebsiella pneumoniae*^4–9^. The inflammatory environment of the gut in IBD increases growth of *Enterobacteriaceae* through the release of bioactive molecules^10,11^, while a dysbiotic microbiota in susceptible mice precipitates inflammation^12^. Particular strains of *E. coli* (Adherent Invasive *E. coli*) and *K. pneumoniae* (Kp2 clade) have been isolated from patients with IBD and cause more severe disease in animal models^13,14^. Additionally, there is evidence that the gut microbiome can serve as a reservoir both for organisms capable of disseminated infection via bacterial translocation and for urinary tract infections^15–18^, of which *K. pneumoniae* is a prominent example.

The gut microbiome is a dense ecosystem, with high levels of microorganism growth and turnover. Resource competition within this environment creates distinct niches, of which different species are capable inhabiting^19^. Two elements necessary for replication and growth of all bacterial species are carbon and nitrogen. Previous work suggests that nitrogen is a limiting resource for growth of organisms in the gut microbiome^20,21^. Nitrogen is essential for anabolic processes including *de novo* synthesis of nucleotides and amino acids. Primary sources for nitrogen in the microbiota are dietary peptides, host-secreted digestive enzymes, glycoproteins from mucus, and the host-produced nitrogenous waste product urea, for which microbes have differing preferences^20,22^. This can in turn affect host physiology, where an intact microbiota increases the daily requirement for host protein intake^23^. Carbon sources are also critical for bacterial survival; in addition to providing carbon for anabolic pathways, these molecules serve as a primary source of ATP. Dietary simple carbohydrates and starches are predominantly metabolized and absorbed by the host in the small intestine, leaving low concentrations available to the microbiota in the colon. Non-digestible complex carbohydrates, known as fiber, fall into the category of microbiota-accessible carbohydrates (MACs). The paucity of simple carbohydrates in the colon favors organisms capable of digesting dietary fibers through glycoside hydrolases (GHs)^24,25^, conveying a significant effect on the microbiome in mouse and human studies^25,26^. While *Bacteroiota* and *Bacillota* can ferment these complex carbohydrates^27^, very little attention has been given to the impact of dietary MACs on *Enterobacteriaceae*, as they do not encode the GH necessary for their utilization^10^. Human gut microbiome response to diet is personalized, as it is dependent upon the pre-existing microbial population and dietary macronutrients^28–30^. Dietary modifications including elimination diet and Mediterranean diets have shown some clinical efficacy in the treatment of IBD, with improvement in dysbiotic signature^31,32^.

We hypothesized that dietary nutrients affect colonization of *Enterobacteriaceae*. Herein, we use bacterial culture, *ex-vivo* studies, and mouse modeling to mechanistically interrogate the results of a previously published human dietary intervention study, dissecting the resource limitation of a prominent member of the *Enterobacteriaceae* family, *K. pneumoniae.* By contrast to the commonly believed importance of nitrogen limitation in the gut, we provide evidence that *Enterobacteriaceae* are not nitrogen limited but rather constrained by simple carbohydrate availability. In contrast, we find that dietary fiber suppresses colonization of *K. pneumoniae* and supports recovery of protective bacterial species after antibiotic depletion in both mouse and human studies. This dietary intervention reduced disease in a model of colitis and remarkably, suppressed systemic dissemination of this human pathogen. Our studies identify dietary factors critical for perpetuation and resolution of dysbiosis in the microbiome through fundamental principles of bacterial metabolism.

## Results

A previously published study from our group examined the impact of dietary fiber on the microbial and metabolomic composition of the gut microbiome in human subjects, referred to as the Food and Resulting Microbial Metabolites (FARMM) study^33^. Healthy human subjects were recruited, of whom 20 normally adhere to a typical American diet (referred to as omnivore diet), and 10 to a vegan diet. The 20 omnivore subjects were randomized to a standardized omnivore diet or a fiber-free exclusive enteral nutrition (EEN) diet, while the vegan subjects remained on a vegan diet (**Figure 1A**). After a five-day dietary phase, all participants were subjected to microbiota depletion via three days of nonabsorbable oral antibiotics (vancomycin and neomycin) together with a one-day polyethylene glycol (PEG) purge. Subsequent recovery of the microbiome was monitored via fecal analysis including shotgun metagenomic sequencing and metabolomics. Previously published results from this study revealed that the diet lacking in fiber, EEN, reduced microbial diversity during microbiome recovery and increased the relative abundance of *Pseudomonadota*^26^, reminiscent of a dysbiotic signature.

**Figure 1.**
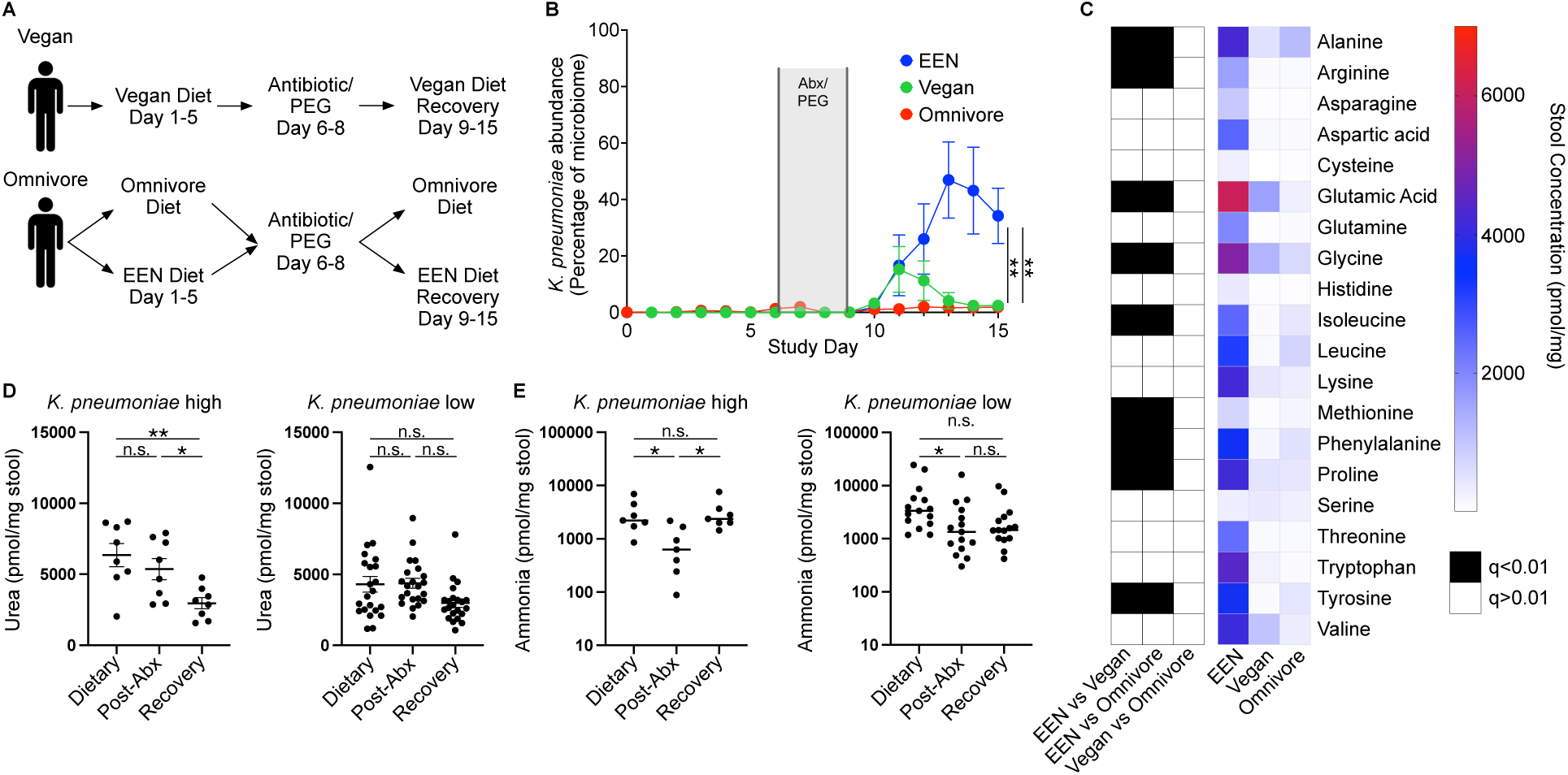
A defined formula diet favors *K. pneumoniae* growth in human subjects. **A**. Diagram of the FARMM study design. Patients were randomized to an FF diet (EEN) or an omnivore diet; a third group remained on a vegan diet throughout the study. Recovery of the microbiome was monitored after antibiotic (abx) and polyethylene glycol (PEG) depletion. **B**. Relative abundance of *K. pneumoniae* as a percentage of the microbiome determined via shotgun metagenomic sequencing, stratified by dietary group. **C**. Heat map of average stool amino acid concentrations during the microbiome recovery phase (Day 14) of the FARMM study, stratified by diet. Black boxes denote statistically significant difference of amino acid concentration between indicated dietary groups. **D and E**. Quantification of stool urea (**D**) and ammonia (**E**) at each phase of the FARMM study, from individuals with high or low relative abundance of K. pneumoniae. Data presented as mean ± SEM, n=10 subjects per dietary group. Results of one-way ANOVA with Tukey correction for multiple comparisons (**B,** comparing EEN vs Omnivore and EEN vs Vegan groups on Day 15**; D, E**) or multiple Mann-Whitney tests with FDR method of Benjamini, Krueger, Yokutieli (**C**), n.s., not significant, *p<0.05, **p<0.01.

Bacterial urease, which metabolizes host-generated urea into ammonia, has been associated with dysbiosis and inflammation in a mouse model of inflammatory bowel disease^34^. Shotgun metagenomic sequencing reveals an increased abundance of the core urease genes in the EEN dietary group relative to the omnivore and vegan groups during microbiome recovery (**Supplemental Figure 1A**). Operon reconstruction reveals that the majority of urease is attributable to *Klebsiella pneumoniae* in the EEN cohort (**Supplemental Figure 1B**). We additionally observed a spike of urease operon genes on the omnivore diet during the early stage of microbiome recovery, which was found to be from *Streptococcus thermophilus* (**Supplemental Figure 1B**), likely attributable to its presence in yogurt in the omnivore diet. Analysis of species abundance revealed that *K. pneumoniae* bloomed in the EEN group, accounting for over 50% of the microbiota, which was not observed in the omnivore and vegan groups (**Figure 1B**).

We hypothesized that colonization of *K. pneumoniae* would alter the fecal nitrogen environment by degrading amino acids and urea into ammonia. During the dietary phase of the study, all 20 proteogenic amino acids are of low abundance (**Supplemental Figure 1C**). Antibiotic treatment increases stool amino acid levels, consistent with previously published study results (**Supplemental Figure 1C**)^26^; however, during microbiome recovery, the EEN dietary group displayed decreased amino acid consumption relative to the omnivore and vegan groups on day 14 of the study (**Figure 1C**). Urea and ammonia quantification revealed that individuals with a high abundance of *K. pneumoniae* had decreased fecal urea and a corresponding increase in ammonia during microbiome recovery, while those with relatively low *K. pneumoniae* levels did not (**Figures 1D and E**). Thus, EEN supported higher levels of *K. pneumoniae* engraftment in human subjects and altered the luminal nitrogen composition of the microbiome.

We hypothesized that urease provides a fitness advantage during intestinal colonization by increasing nitrogen availability. We generated an *in vitro* model system to explore the role of urea in a complex gut microbial community. A bioreactor system was inoculated with a human fecal sample into two complex media: 1. Brain Heart Infusion (BHI) medium, in which the primary carbohydrate is glucose, or 2. SHIME medium, which contains complex glycans^35^. The low concentration of urea in both media provided the opportunity to directly test the impact of urea on an intestinal microbial community. Samples were collected every 24 hours and subjected to shotgun metagenomic sequencing with relative species and gene abundances calculated. We found that in BHI, the addition of urea significantly increased the abundance of the urease-positive organisms *K. pneumoniae* and *Citrobacter freundii* (**Supplemental Figures 2A and B**). The abundance of urease genes accordingly increased with urea supplementation (**Supplemental Figure 2C**). These relationships were substantially diminished in SHIME media (**Supplemental Figures 2A-C**), supporting the theory that urease provides a growth advantage in the presence of urea through increased nitrogen availability only when a simple carbohydrate is available for use by *Enterobacteriaceae*.

To rigorously test the hypothesis that nitrogen is growth limiting in the gut for *K. pneumoniae*, we generated two deletion mutants: the urease operon (Δurease) and *ntrC* (Δ*ntrC*), which encodes a key regulator of the nitrogen scavenging Ntr system^36^. While the WT strain grew *in vitro* well in an ammonia-limited environment with urea as an available nitrogen source, the Δ urease and Δ*ntrC* strains displayed delayed or limited growth (**Supplemental Figures 2D and E**). In an ammonia-rich environment, urea had no effect on the three strains. When tested on a range of nitrogen-containing compounds, the Δ*ntrC* strain was significantly restricted in its growth relative to the WT strain (**Supplemental Figure 3A**). Thus, we find that under nitrogen-limited environments *in vitro*, the *K. pneumoniae* has a broad capacity for nitrogen assimilation, dependent on urease and *ntrC*.

To directly test if restricting the nitrogen resources available to *K. pneumoniae* reduces colonization capacity in the gut, mice were colonized with WT, Δurease, or Δ*ntrC K. pneumoniae* after antibiotic pre-treatment on a standard mouse chow (**Supplemental Figures 3C and D**). Surprisingly, we found no colonization defect for the mutant strains. This led us to speculate that under normal dietary conditions, the gut is abundant in available nitrogen; previous work demonstrated that limitation of dietary protein significantly reduced ammonia and urea in the mouse intestine^37^. However, we find that urease and *ntrC* are also dispensable for colonization of mice on a low-protein diet (**Supplemental Figures 3E and F**). The absence of complex carbohydrates similarly had no impact on colonization of Δurease *K. pneumoniae* (**Supplemental Figures 3G and H**). We confirmed that urease did alter the intestinal nitrogen environment, increasing ammonia levels with corresponding urea consumption (**Supplemental Figure 3I-N**).

To rule out the effect of cross-feeding from other members of the microbiota on colonization of urease-null *K. pneumoniae*, we colonized germ-free (GF) mice with WT or Δurease *K. pneumoniae*. Intestinal colonization of *K. pneumoniae* was independent of urease expression (**Figure 2A**). We found that, prior to gavage, fecal ammonia levels were remarkably low (**Figure 2B**), similar to that of microbiome depleted conventionally raised (CR) mice (**Supplemental Figure 3J**). Colonization with WT *K. pneumoniae* resulted in ∼10-fold increase in fecal ammonia levels; this was partially urease-dependent as colonization with Δurease *K. pneumoniae* resulted in significantly less ammonia production than the WT strain but remained ∼4-fold above the GF baseline. WT *K. pneumoniae* colonization caused decreased fecal urea levels relative to the urease-deficient strain, demonstrating urea hydrolysis as a primary source of ammonia production (**Figure 2C**). Fecal amino acid analysis reveals a significant decrease in the concentration of several amino acids after colonization with WT strain (**Figure 2D**), likely due to consumption by *K. pneumoniae*. Combined with the production of gut ammonia by the Δurease *K. pneumoniae* strain, this reduction of fecal amino acids suggests the utilization of amino acids as a carbon source via deamination, thereby leading to the release of ammonia into the gut environment.

**Figure 2.**
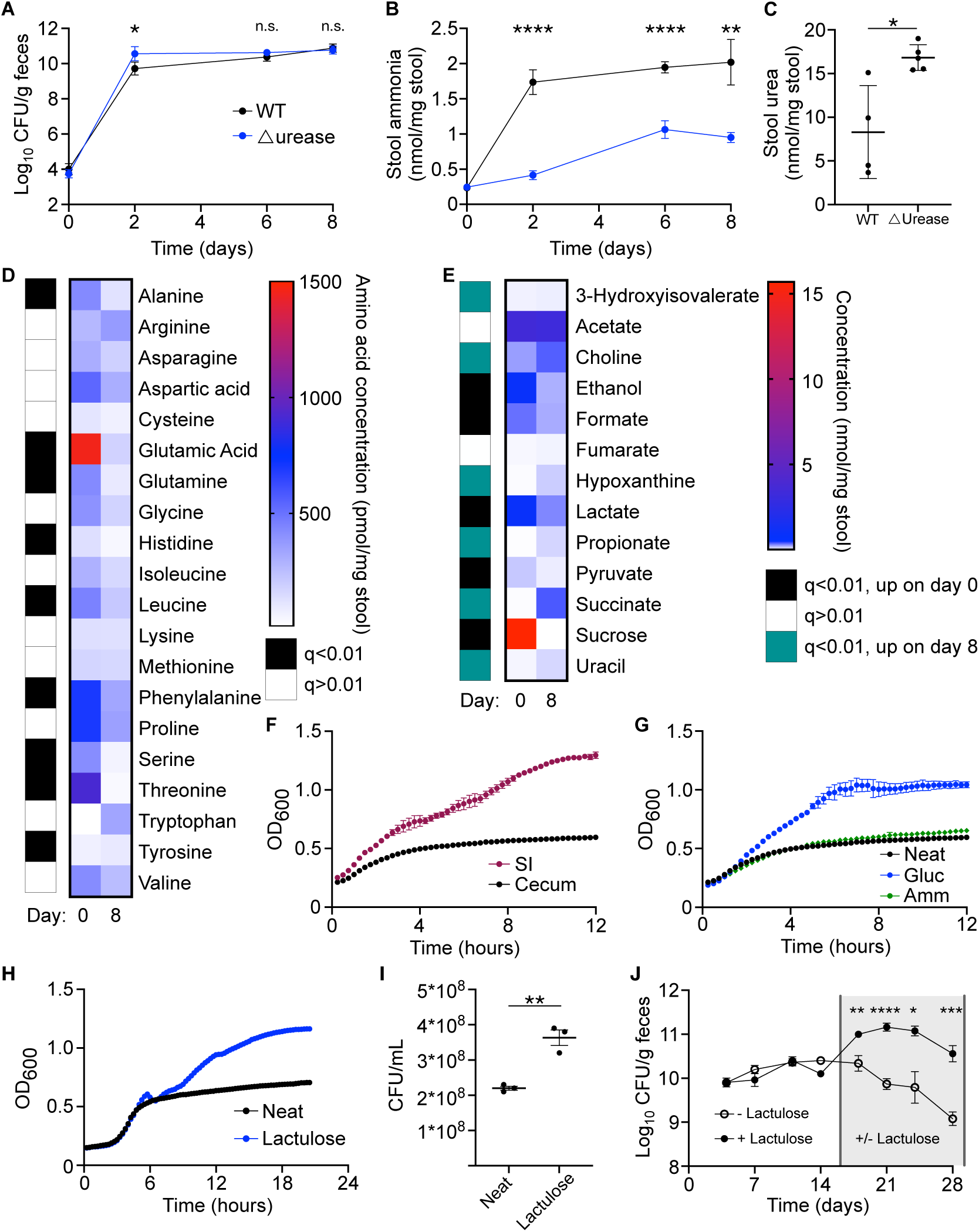
*K. pneumoniae* colonization causes increased fecal ammonia associated with consumption of urea, amino acids, and carbon sources. **A-E**. Germ-free mice were colonized with WT or Δurease *K. pneumoniae* and serial stool collections performed throughout the 1-week study. Fecal CFU (**A**) and stool ammonia (**B**) were monitored for 1 week after colonization. Fecal urea (**C**) was tested at day 8. Stool amino acid levels (**D**) and metabolites (**E**) were quantified from stool before (day 0) and after (day 8) WT K. *pneumoniae* colonization. **F.** Growth of WT *K. pneumoniae* in small intestine (SI) or cecal extract of mice monitored via OD_600_. **G and H**. Growth in cecal extract supplemented with ammonia or glucose (**G**), or lactulose (**H and I**) quantified by OD_600_ and CFU (**I**). **J**. Mice colonized with *K. pneumoniae* were subsequently treated with lactulose in the drinking water or water-only control. Data presented as mean ± SEM (**A and J**) or mean ± SD (**B, C**, **and F-I**); n=4-5 mice per group (**A-E, J**) or n=3 wells per group (**F-I**). Data represents combined results from two independent experiments (**A-E**) or are a single experiment representative of three independent experiments (**F-J**). Results of one-way ANOVA with Bonferroni correction for multiple comparisons, n.s., not significant, *p<0.05, **p<0.01, ***p<0.001, ****p<0.0001.

The observed pattern of amino acid consumption and ammonia production after *K. pneumoniae* colonization indicated that carbon may be the primary limited resource for colonization of *K. pneumoniae* in the gut. Through *in vitro* testing, we found that *K. pneumoniae* has a strong preference for simple carbohydrates, in addition to select amino acids, and citric-acid cycle intermediates (**Supplemental Figure 3B**), while complex carbohydrates including inulin and dextrin are not utilized. Proton NMR quantification of fecal samples before and after colonization of GF mice identified several metabolites which were significantly reduced in concentration after *K. pneumoniae* colonization, including sucrose, pyruvate, and lactate (**Figure 2E**), all of which serve as carbon sources for *K. pneumoniae* in culture (**Supplemental Figure 3B**). These data support that *K. pneumoniae* is carbon limited in the intestinal environment, and that alternative sources of nitrogen are unnecessary for colonization.

The mammalian small intestine absorbs most simple carbohydrates and amino acids, supporting the hypothesis that *K. pneumoniae* is limited by carbon sources available in the colon. We generated an *ex-vivo* model in which the small intestinal or cecal content of microbiota-depleted CR mice were extracted, sterilized, and used as culture media for growth of *K. pneumoniae*. Growth in the small intestinal extract was greater than that in the cecal content, suggesting that host-absorbed nutrients are important for intestinal growth of *K. pneumoniae*. Addition of the readily useable nitrogen source, ammonia, to cecal extract had no effect on growth of *K. pneumoniae*, however, glucose supplementation increased maximum density and growth rate (**Figures 2F and G**) to mirror that of the SI extract.

One simple carbohydrate that is not absorbed or metabolized by the host is lactulose, an artificial disaccharide of galactose and fructose. We found that lactulose can be utilized as a sole-carbon source in minimal media for the tested strain of *K. pneumoniae* (**Supplemental Figure 3B**). We confirmed that addition of lactulose to *ex-vivo* cecal extract increased growth of *K. pneumoniae* measured by optical density and CFU (**Figures 2H and I**). Mice were provided lactulose in drinking water after colonization with WT *K. pneumoniae*. Compared to water control, lactulose increased *K. pneumoniae* colonization by 10-fold (**Figure 2J**). In total, these data provide strong evidence that colonization of *K. pneumoniae* in the gut microbiome is carbon restricted and that nitrogen is abundant in the intestinal environment.

While available simple carbohydrates support the growth of *K. pneumoniae* in the gut, the majority of carbohydrates in the distal gut are complex carbohydrates, or fiber, that are not absorbed by the host. We hypothesized that these complex carbohydrates, referred to as dietary fiber, may play a role in colonization resistance against *K. pneumoniae*. Indeed, results of the FARMM study demonstrate that the EEN diet, which is deficient in fiber, increased colonization of *K. pneumoniae*. To test this hypothesis, we generated matched fiber-free (FF) and high-fiber (HF) mouse chows; the HF chow contained a well-characterized pea fiber^38^. Groups of CR mice were provided FF or HF chow *ad libitum*, treated with the same oral antibiotic regimen provided to the FARMM subjects for three days to deplete the microbiota, and gavaged with *K. pneumoniae*. As expected, we find that antibiotic treatment significantly reduces microbiome diversity when compared via shotgun sequencing (**Figure 3A**). During recovery, microbiome diversity was dependent upon dietary fiber, where the mice provided a HF diet returned to baseline diversity by 4 weeks and FF diet precluded such recovery. This data mirrors that of the FARMM study, wherein human subjects on a fiber-free EEN diet did not recover to baseline diversity after microbiome depletion. PCA analysis confirms that while antibiotic treatment significantly affects the community structure regardless of diet, the community on the FF diet remains persistently altered from baseline and the HF group (**Supplemental Figure 4A**). We hypothesized that fiber would enable asymmetric recovery of organisms encoding glycoside hydrolases (GHs) necessary for complex glycan metabolism. Indeed, while mice on a HF diet had increased levels of GHs enabling metabolism of plant-based carbohydrates, these were decreased in mice fed an FF diet (**Figure 3B**). In contrast, GHs for simple carbohydrates and animal glycans were disproportionately increased in the FF diet group, reminiscent of results from the FARMM study and consistent with the ability of a FF diet to select for a mucolytic community^39^.

**Figure 3.**
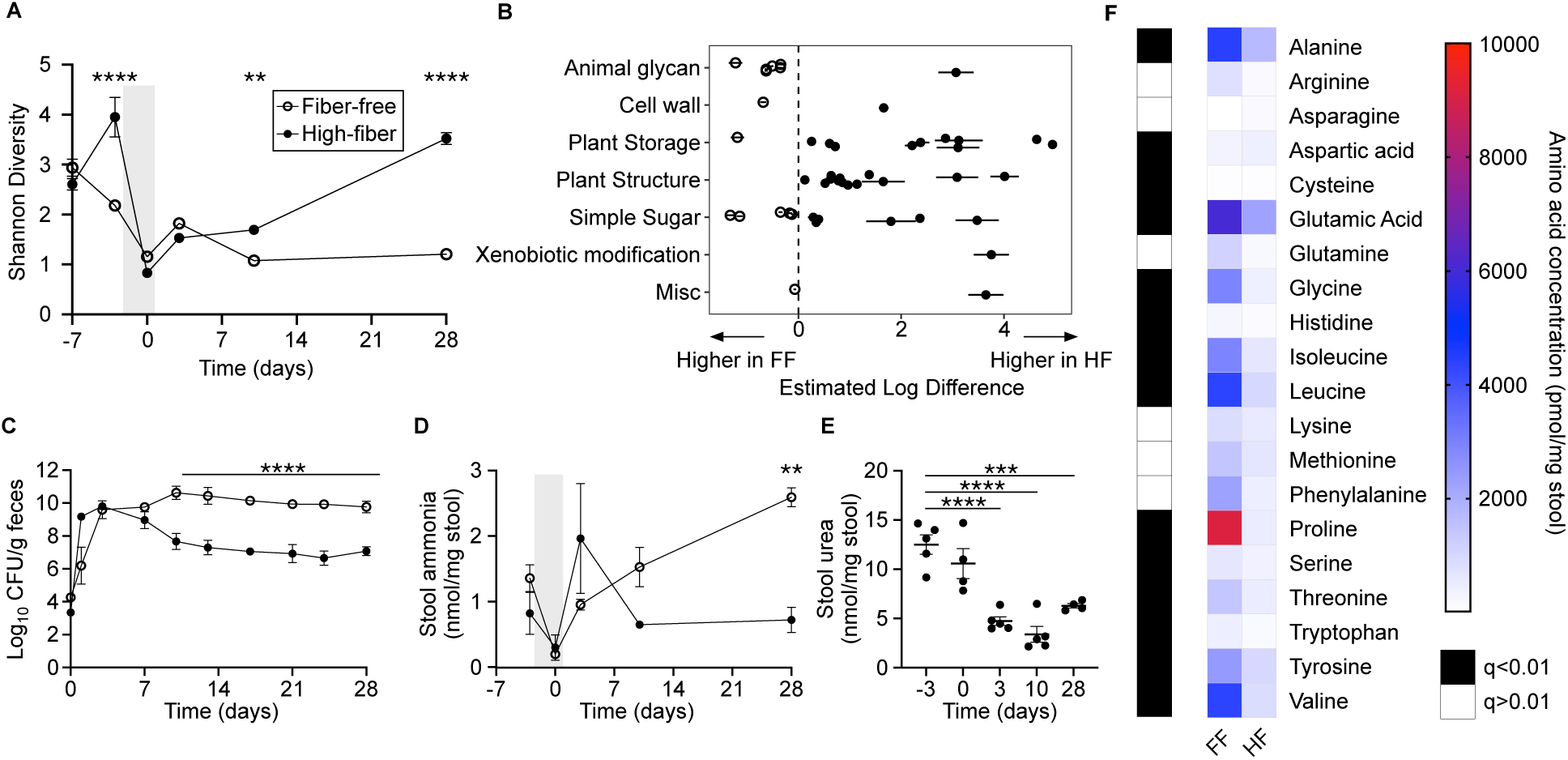
Complex carbohydrates increase microbiome diversity and reduce *K. pneumoniae* colonization after antibiotic depletion. **A-F**. Mice were provided a fiber-free (FF) or high-fiber (HF) diet (starting at day -7), treated with oral antibiotics (day -3 to 0, grey shading), and gavaged with *K. pneumoniae* (day 0). Serial stool samples were subjected to analysis as follows: **A.** Shannon alpha-diversity of stool microbiome from mice provided a fiber-free (FF) or high-fiber (HF) diet including antibiotic treatment period (grey shading) and recovery, as determined by shotgun metagenomic sequencing. **B.** Glycoside hydrolase (GH) genes with significantly different abundances between FF and FH diets four-weeks after gavage with K. pneumoniae grouped by substrate type. Open circles represent genes with higher levels in FF diet and closed circles represent genes with higher levels in HF diet. **C.** *K. pneumoniae* fecal CFU of mice on FF (open circles) or HF (closed circles) diets measured four weeks after *K. pneumoniae* gavage. **D**. Stool ammonia was quantified before and after antibiotic treatment (grey shading) and *K. pneumoniae* gavage from mice subjected to FF (open circles) or HF (closed circles) diets. **E.** Stool urea levels were quantified from mice provided a FF diet. **F**. Heat map of amino acid concentrations from mice on an FF or HF diet after colonization with *K. pneumoniae*. Data are presented as mean ± SD (**A-E**), n= 5 mice per group. Data representative of two to three independent experiments (**C-F**). Results of one-way ANOVA with Bonferroni correction for multiple comparisons (**A, C-E**), n.s. not significant, **p<0.01, ***p<0.001, ****p<0.0001. Results of multiple Mann-Whitney tests with FDR method of Benjamini, Krueger, Yokutieli (**F**), *q<0.02.

After gavage with *K. pneumoniae*, we found 1000-fold higher level of colonization in the FF group compared with the HF group (**Figure 3C**). Intriguingly, this difference only manifested after one week, corresponding with divergence in microbiome diversity between groups. Species analysis of the microbiome through shotgun metagenomics identified several organisms that were permanently lost on the FF diet, the recovery of which on a HF diet anti-correlated with *K. pneumoniae* (**Supplemental Figures 4B and C**). Several members of the microbiota including *Lactobacillus johnsonii*, *Bifidobacterium pseudolongum*, and a *Lachnoclostridium* species demonstrated a strong correlation with resistance to *K. pneumoniae* colonization (**Supplemental Figures 4B and C**), suggesting that these species may play a causative role in the effect of this dietary intervention. Indeed, analysis of metagenomic data from the FARMM trial found a similar pattern, in which *Lactobacillus* and *Bifidobacterium* species were at lower levels in the EEN dietary group compared with fiber-containing diets (**Supplemental Figure 4E**).

Examination of the nitrogen composition of the gut demonstrates that stool ammonia is increased in the FF group, corresponding with higher levels of *K. pneumoniae* and associated with urea depletion over the course of the study (**Figures 3D and E**). By contrast, we did not find such a relationship in the HF diet group (**Figures 3D and E; Supplemental Figure 4F**) similar to the results of the FARMM study (**Figures 1D and E**). Moreover, we observed higher levels of stool amino acids in the FF group relative to the HF group during microbiome recovery (**Figure 3F**), suggesting that organisms abundant on the HF diet preferentially consume amino acids. In summary, we have replicated several key features of the human subject FARMM study through mouse modeling by removing dietary fiber, including altered microbiome diversity, GH abundance, *K. pneumoniae* colonization, and nitrogen composition. Our results demonstrate that complex dietary carbohydrates play a key role in colonization resistance against *K. pneumoniae*.

One population for whom a fiber-free diet is routinely prescribed are critically ill patients receiving EEN. Bacterial translocation (BT) from gut barrier dysfunction has long been speculated as a source of infection in these patients. However, the dietary factors predisposing to dissemination of *Enterobacteriaceae* have not been explored. To test the impact of dietary fiber on BT, mice were colonized with *K. pneumoniae* on an FF or HF diet after antibiotic pre-treatment. After stable engraftment, mice were treated with dextran sodium sulfate (DSS) to induce intestinal inflammation and disrupt the epithelial barrier. Consistent with previous studies, we found that disease activity, weight loss, and colon length were improved by the addition of dietary fiber (**Figures 4A-C**)^40^. After four days of treatment, mice were euthanized and *K. pneumoniae* quantified from sections of the intestinal tract, liver, and blood. Dietary fiber not only reduced colonization of *K. pneumoniae* in the colon, but also in the proximal and distal small intestine (**Figures 4D-F**). We observed that fiber prevented dissemination of *K. pneumoniae* to the liver and blood, while mice treated with an FF diet had greater levels of bacteremia and liver infection (**Figures 4G and H**). In total, our data unveil a novel role for dietary carbohydrates on colonization of *K. pneumoniae* in the gut and provide a possible treatment to reduce translocation of this human pathogen.

**Figure 4.**
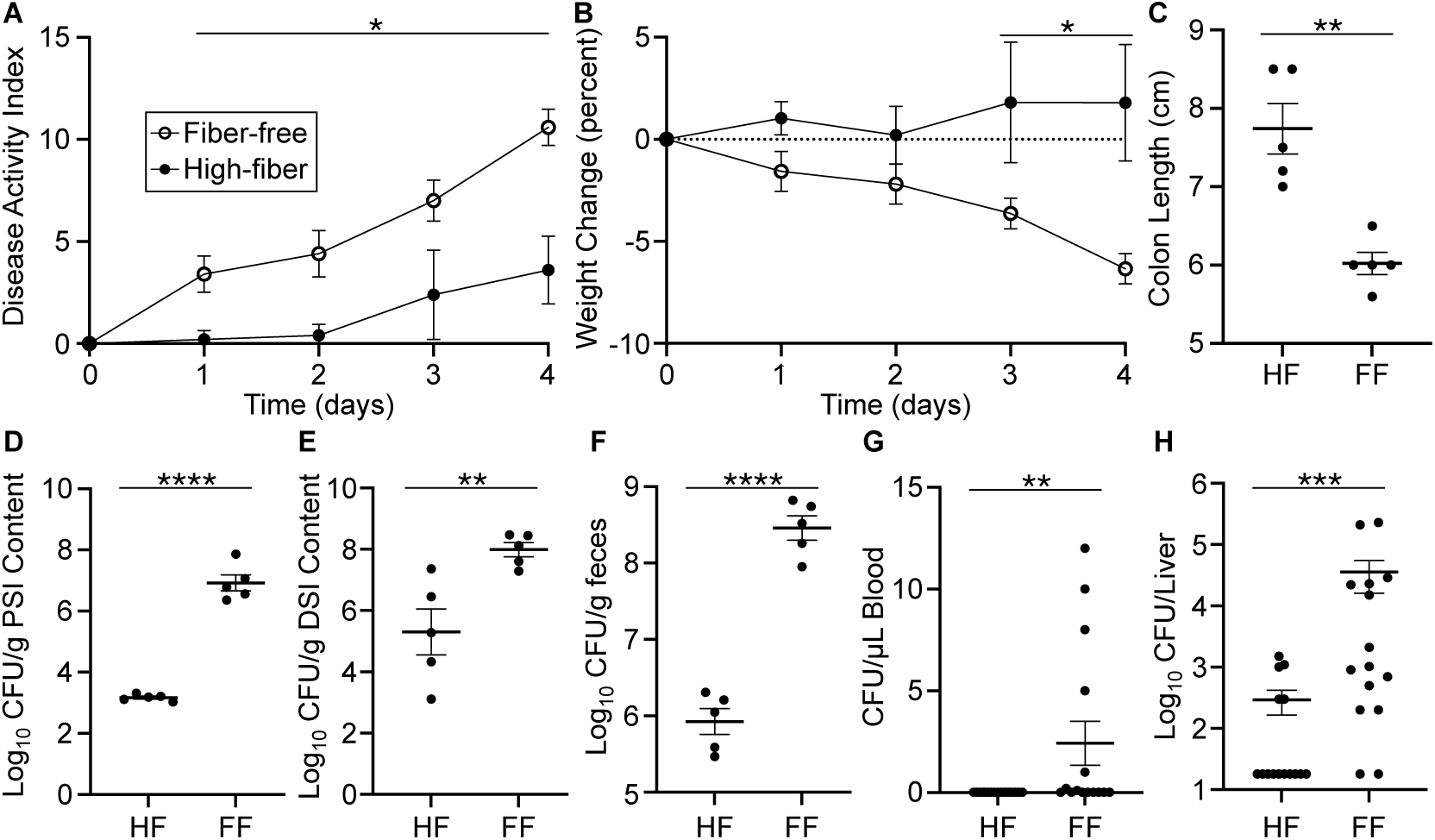
High fiber diet protects against *K. pneumoniae* dissemination from the microbiota. **A and B**. Mice were placed on a HF or FF diet and colonized with WT *K. pneumoniae* after antibiotic pretreatment. Subsequent treatment with 5% DSS in the drinking water was performed, with monitoring of disease activity (**A**) and weight change (**B**). **C-H.** After 4 days of treatment, mice were euthanized. Colon length measured at this time point (**C**) and CFU was quantified in the proximal small intestine (**D**), distal small intestine (**E**), feces (**F**), blood (**G**), and liver (**H**). Data are presented as mean ± SD, n= 5 mice per group. Results representative of three independent experiments (**A-F**) or a combination of three independent experiments (**G and H**). Results of multiple Mann-Whitney tests with FDR method of Benjamini, Krueger, Yokutieli (**A and B**), *q<0.001. Results of student’s t-test (**C-F**) or Mann-Whitney test (**G and H**), **p<0.01, ***p<0.001, ****p<0.0001.

## Discussion

Dysbiosis is an important feature of many chronic diseases, including IBD, and involves the outgrowth of *Enterobacteriaceae* in the gut. *K. pneumoniae* and other *Enterobacteriaceae* are not only thought to worsen IBD but also serve as infectious human pathogens in bacteremia, cholangitis, urinary tract infections, and pneumonia. Identifying dietary components contributing to the formation and resolution of dysbiosis is therefore critical. We find that diet can have a direct effect on colonization of *K. pneumoniae* by serving as a nutrient source and an indirect effect via the microbiota. Understanding the dietary resources necessary for colonization of harmful bacterial strains could serve as a critical tool in targeted prebiotic therapies.

Previous literature suggests that nitrogen is a limiting resource for bacterial growth in the mammalian gut^20,21^. Urea is a host-produced nitrogenous waste product which is in turn excreted into the GI tract and has long been speculated as an important source of nitrogen for the microbiome, accessed via bacterial urease^27,41^. Indeed, recent literature shows that bacterial urease is important for cross-feeding of nitrogen to non-urease encoding species^22^. Our human subject and bioreactor data suggested that urease may be advantageous for the growth of *K. pneumoniae* in microbiome recovery, wherein levels of the readily accessible nitrogen source, ammonia, is less abundant. However, after rigorous testing *in vitro*, *ex-vivo*, and *in vivo*, we find that nitrogen is not a limiting resource for the colonization of *K. pneumoniae*. Even under nitrogen-starved conditions, bacterial urease and the nitrogen scavenging system, Ntr, is dispensable for growth of *K. pneumoniae*. This suggests that nitrogen is abundant in the mouse intestinal tract and that other dietary factors are more central for the colonization of this human pathogen.

Regardless, *K. pneumoniae* colonization has a significant impact on the nitrogen environment of the intestinal tract. Only select amino acids are consumed after *K. pneumoniae* mono-association of GF mice, including serine and threonine, which are most likely being used as carbon sources through deamination. *K. pneumoniae* colonized GF mice maintain higher levels of many amino acids compared to CR mice or humans with intact microbiota; other members of the community, particularly *Clostridia*, are capable of Stickland fermentation and thus have a broader accessibility to energy extraction from amino acids^42^. Interestingly, we find that luminal ammonia, which serves as a preferred nitrogen source for most bacterial species, is dependent on the presence of the gut microbiome, where feces of GF mice and antibiotic-treated mice and humans contain substantially lower levels. Our work demonstrates that while urease is a factor in ammonia production during colonization of GF mice, ammonia is also produced from other sources, likely including amino acid deamination. It is possible that other bacterial species in the gut microbiome remain nitrogen limited in the gut, including organisms in the phyla *Bacteroidota* and *Bacillota*, as many are capable of consuming complex carbohydrates, thus shifting the ratio of available carbon to nitrogen. Indeed, it has been speculated that *Bacteroides* species are dependent on ammonia as their core nitrogen source^27^; our data suggests that ammonia is abundant in the presence of a gut microbial community, with contributions from urea degradation and amino acid deamination. Further studies will be needed to characterize the available luminal nitrogen sources for these phyla.

We find that carbon sources are a major limiting factor for growth of *K. pneumoniae* in the gut. The host absorbs most simple carbohydrates in the small intestine, leaving a relative paucity of dietary simple carbohydrates available to the microbiota in the colon. Previous work focused on *E. coli* has identified multiple carbohydrate utilization genes important for competitive colonization of the GI tract, including gluconate, mannose, fucose, and ribose^43^, thought to be liberated from host mucus polysaccharides^44^. While *K. pneumoniae* carbohydrate consumption in the intestine is less well studied, recent work demonstrated that fucose utilization is important for intestinal colonization^45^. The role of dietary carbohydrates in colonization of *Enterobacteriaceae* has previously not been well characterized. Our *ex-vivo* modeling reveals that growth in SI luminal extract was greater than that of the cecum, the effect of which was diminished with the addition of a simple carbohydrate to the cecal material. Moreover, supplementation with a non-absorbed carbohydrate, lactulose, in a mouse model increased colonization of *K. pneumoniae*, consistent with the hypothesis that carbohydrates are a major limiting resource for colonization. Host absorption of simple carbohydrates in the small intestine may thus play a role in colonic colonization resistance against pathobionts in this family.

Conversely, we find that complex carbohydrates found in dietary fiber suppresses colonization of *K. pneumoniae* both in humans and in mice correlating with increased microbiome diversity. Consistent with the FARMM study, dietary fiber enriched for plant-based GH abundance, whereas a fiber-free diet favored GHs for simple carbohydrates and animal glycans^26^. Short chain fatty acids (SCFAs), often produced in the intestine from dietary fiber, has also been shown to suppress *K. pneumoniae* growth via intracellular acidification^46^. This suggests that fiber may have two roles in inhibiting *Enterobacteriaceae* colonization: directly through SCFA production and via supporting the recovery of competing organisms encoding plant saccharide GHs. *Clostridioides difficile* colonization provides some analogy, as it also relies on both depletion of microbiome diversity and decreased SCFA production^47^. Further studies will be necessary to define the causal strain(s) and the essential GHs for this phenotype.

The suppression of *K. pneumoniae* colonization in the microbiome would have significant disease implications. For patients with IBD, reducing *K. pneumoniae* colonization improved host inflammatory response^14^. Moreover, disseminated *K. pneumoniae* infections, in part, derive from the gut microbiome^48^. Indeed, we find that disruption of the intestinal barrier with DSS leads to dissemination of *K. pneumoniae* to the liver and bloodstream, which is nearly eliminated by reduction of colonization with dietary fiber. Lack of fiber in the diet has separately been shown to increase the severity of DSS and infectious colitis, and thus the effect seen is likely multifactorial^40,49^.

Our studies reveal the critical importance of dietary carbohydrates in colonization of *K. pneumoniae* in the mammalian gut. This study may have direct relevance to human disease; patients hospitalized in the intensive care unit often require EEN, which is typically given without fiber supplementation. The outgrowth of *Enterobacteriaceae* in this setting due to poor microbiome diversity may predispose these patients to disseminated infection. Resolution of dysbiosis through rational prebiotic and probiotic therapies would have a range of therapeutic applications. Understanding the limiting metabolic resource(s) for these disease-associated organisms and the impact of diet on their colonization is a key step in designing such treatment strategies.

## Resource Availability

Further information and requests for resources should be directed to Gary D. Wu

## Materials Availability

Reagents and strains generated from this study will be made available upon reasonable request with appropriate materials transfer agreement.

## KEY RESOURCES TABLE

| REAGENT or RESOURCE | SOURCE | IDENTIFIER |
| --- | --- | --- |
| **Bacterial and virus strains** |  |  |
| *Klebsiella pneumoniae* MGH 78578 | ATCC | 700721 |
| *K. pneumoniae* Δurease | This study | NA |
| *K. pneumoniae* ΔntrC | This study | NA |
| **Biological samples** |  |  |
| Human fecal samples | FARMM study | NA |
| **Chemicals, peptides, and recombinant proteins** |  |  |
| Kanamycin Sulfate | Fisher | BP906 |
| Vancomycin Hydrochloride | Fresenius Kabi | NDC 63323-314-61 |
| Neomycin Trisulfate Salt Hydrate | Sigma | N6386 |
| Hygromycin B | Corning | 30-240-CR |
| Apramycin Sulfate | Alfa Aesar | J66616 |
| Polyethylene Glycol 3350 | Miralax | NA |
| Dextran Sulfate, Sodium Salt | Fisher | BP1585 |
| Lactulose | Thermo Scientific | J60160.22 |
| Phenotype Microarrays | Biolog | PM1, PM2A, PM3B |
| **Critical commercial assays** |  |  |
| Quantichrom Urea Assay Kit | BioAssay Systems | 76237-110 |
| **Deposited data** |  |  |
| FARMM Shotgun Metagenomics | FARMM study | PRJNA675301 |
| **Experimental models: Organisms/strains** |  |  |
| C57Bl/6J | Jackson Laboratory | NA |
| **Oligonucleotides** |  |  |
| See Supplemental Table 1 |  |  |
| **Plasmids** |  |  |
| pMDIAI | Addgene | 51655 |
| pFLP-hyg | Huang,T *et al.* | NA |
| pACBSR-hyg | Huang,T *et al.*<sup>50</sup> | NA |

| Software and algorithms |  |  |
| --- | --- | --- |
| Prism 9.5.1 | GraphPad | NA |

## EXPERIMENTAL MODEL AND SUBJECT DETAILS

### Human Subjects

Stocked fecal samples from a previously reported human study called Food and Resulting Microbial Metabolites (FARMM) were tested where noted^26^. In brief, the FARMM study consisted of 30 healthy human subjects, 10 of whom were vegan at baseline, 20 of whom had a typical Western (omnivore) diet. Vegan subjects maintained an outpatient vegan diet during the study. The omnivore group were randomized to a standardized omnivore diet or exclusive enteral nutrition (EEN) diet, consisting of Modulen® IBD. The omnivore standardized diet was designed to have similar composition, excepting the lack of dietary fiber in EEN. All subjects underwent three days of oral vancomycin 500mg and neomycin 1000mg every 6 hours on days 6, 7, and 8 of the study. All subjects were also given a bowel purge consisting of 4L polyethylene glycol purgative (GoLytely®). The University of Pennsylvania Institutional Review Board (IRB) approved the protocol and considered it exempt from clinical trial registration requirement.

### Mice

All animal studies were conducted in accordance with ethical regulations under protocols approved by the University of Pennsylvania Institutional Animal Care and Use and Biosafety Committees. SPF C57Bl/6J female mice were purchased from Jackson Laboratories at 8 weeks of age. Mice were housed under standard lighting cycle conditions (12 hours on/12 hours off) and provided acidified water. There was no investigator blinding for these studies and no animals were excluded from analysis.

### Mouse *K. pneumoniae* Colonization Model

All CR mice were specific pathogen free (SPF) C57Bl/6J strain females acquired from Jackson Laboratories at 8 weeks of age. For colonization with *K. pneumoniae*, mice were treated with the antibiotics vancomycin (2.5g/L) and neomycin (5g/L) with aspartame (25g/L) in the drinking water provided *ad libitum* for 72 hours. Mice were then orogastrically gavaged with 10^8^ CFU of *K. pneumoniae* in 100µL, the inoculum of which was generated from aerobically grown overnight culture in LB media, washed and diluted 1:10 in sterile PBS. Fecal samples for CFU analysis were collected, homogenized in PBS, serially diluted, and spot plated on LB agar supplemented with kanamycin and vancomycin. Plates were incubated for 12 hours at 37°C, colonies counted, and CFU calculated per weight of fecal sample. Fecal samples collected for metabolomics analysis were flash frozen in dry ice and stored at -80℃. Diets were irradiated and provided *ad libitum*; the following diets were used in the indicated experiments: standard diet (Research Diets, AIN-76A), fiber-free diet (TestDiet 5Z6G), high fiber diet (TestDiet 5Z6L; containing Vitacel Pea Fiber EF-100 from J. Rettenmaier, USA), and low protein diet (Research Diets, D08092201).

For lactulose treatment experiments, mice were colonized with *K. pneumoniae* and subsequently provided 5% lactulose w/v in drinking water *ad libitum* after two weeks of colonization.

### Mouse DSS and Dissemination Model

For DSS colitis model, mice were treated with 5% DSS in the drinking water. Mice were monitored for disease activity, as per previously published protocols^34^. After 4 days of treatment, mice were euthanized, and liver and blood harvested. Luminal content was collected at the time of euthanasia from the proximal 10cm of small intestinal, distal 10cm of small intestine, and fecal pellets. Colon length was measured at the time of euthanasia. Liver samples and intestinal luminal contents were homogenized and all samples were serially diluted and spot plated for CFU on LB with kanamycin and vancomycin. CFU were calculated per weight or per liver, as indicated.

### Germ Free Mouse Colonization Model

For germ-free animal experiments, C57Bl/6J female mice were raised and maintained in sterile conditions in the University of Pennsylvania Gnotobiotic Facility. For colonization, mice were orogastrically gavaged at 8-weeks of age with indicated *K. pneumoniae* strains. Fecal samples were harvested before and after gavage at indicated time points and subjected to noted CFU and metabolomics analysis.

### Bioreactor model

Eppendorf (Hamburg, Germany) Bioflo 320 bioreactors were assembled according to the manufacturer’s specifications. During the experiment, all bioreactors were maintained at a temperature of 37 °C with agitation set to 100 rpm. The pH was kept at 7 ± 0.1, using 1 M NaOH and CO_2_ gas. There was a constant sparging of gas at a rate of 1L/min, with the ratio of N_2_:CO_2_ dependent on the pH control. The initial volume of the bioreactors was 750 mL of either Brain Heart Infusion (BHI) broth) or a 70:30 ratio of Adult M-SHIME® growth medium:Pancreatic Juice. Adult M-SHIME® growth medium with starch was purchased from ProDigest (Ghent, Belgium) and consists of arabinogalactan (1.2 g/L), pectin (2 g/L), xylan (0.5 g/L), glucose (0.4 g/L), mucin (2 g/L), and starch (4 g/L), as well as yeast extract (3 g/L) and peptone (1 g/L) which provide nitrogen and trace nutrients [PMID 30016326]. Pancreatic juice was made fresh every 2-3 days and contained 12.5 g NaHCO_3_ (Sigma-Aldrich, St. Louis, MO), 6 g oxgall bile (Becton-Dickinson, Franklin Lakes,NJ) and 0.9 g pancreatin (Sigma-Aldrich, St. Louis, MO)^35^. The bioreactors were connected using silicon tubing (Cole Parmer, Vernon Hills, Il) to a supply of medium and pancreatic juice to provide nutrition and biliary-pancreatic enzymes and to a bard urinary drainage bag (Becton, Dickinson and Company, Franklin Lakes, NJ) to collect waste. For inoculation, fecal samples were suspended at 10% wt/v in phosphate buffer within an anaerobic chamber. Buffer consisted of 0.8 g K_2_HPO_4_, 6.8 g KH_2_PO_4_, 0.1 g Sodium thioglycolate, 15 mg sodium thionite per liter with pH adjusted to 7. At time of inoculation, 10 mL of fluid was removed from the bioreactor and 10 mL of the resuspended inoculums added. The bioreactors were grown overnight, 16h, with temperature, pH, agitation, and gas flow. Following overnight growth, the bioreactors were maintained with daily feeding cycles in which the bioreactor was provided fresh medium and pancreatic juice. Every 8 h, the volume of the bioreactor was reduced to 600 mL and 150 mL of either fresh BHI or a 70:30 Adult M-SHIME® growth medium:Pancreatic Juice mixture was added. Residence time was 40 h.

Reactors were inoculated with a baseline fecal samples from a healthy human subject enrolled in a previously described dietary intervention study^26^. Following a 14-day period at baseline conditions, urea was supplemented to the feed at a concentration of 10 mM for an additional 14-day period. Samples were collected in the morning prior to the beginning of the feeding cycle.

### Ex vivo model

C57Bl/6J female mice 8 weeks of age were obtained from Jackson Laboratories and treated with antibiotics as previously noted for 72 and the final 24 hours with 10% polyethylene glycol in the drinking water. Regular water was then resumed for 24 hours. Mice were then euthanized and small intestinal or cecal luminal content was harvested, weighed, and resuspended in M9 minimal media without ammonia or glucose supplementation at a ratio of 4mL:1g material.

Samples were homogenized for 10 minutes and centrifuged at 10,000 x g for 10 minutes twice. Supernatant was filter sterilized through syringe driven 0.45µm PTFE filter. For aerobic experiments, overnight aerobic culture of *K. pneumoniae* grown in LB was washed in PBS, diluted 1:1000, and inoculated into 200µL of SI or cecal extract in 96 well plates. For anaerobic experiments, SI or cecal extracts were allowed to equilibrate in anaerobic chamber for 24 hours and inoculated with PBS-washed anaerobic overnight culture of *K. pneumoniae*, diluted 1:1000. Growth was monitored for 24 hours via OD_600_ in BioTek Epoch2 plate reader, incubated at 37°C. CFU was obtained via serial dilution and spot plating on LB agar with kanamycin and vancomycin after 24 hours of growth.

## METHOD DETAILS

### Bacterial strains, culture conditions, and antibiotics

*K. pneumoniae* strain used was MGH 78578, as noted in Supplemental Table 1. *K. pneumoniae* was grown in LB (miller) medium aerobically at 37°C unless otherwise noted. M9 minimal media was used with carbohydrate or nitrogen supplementation as follows: 0.1% ammonia, 0.2% glucose, or 0.2% lactulose where noted. Antibiotics used were as follows: kanamycin, apramycin, vancomycin, and hygromycin at concentrations of 25, 100, 8, and 50µg/mL, respectively.

### Strain Construction

Deletion of indicated genes in the MGH 78578 *K. pneumoniae* strain was performed via Recombineering through replacement of the targeted with a FRT-apramycin-FRT construct and subsequent excision, as previously described^51^. In brief, the pMDIAI plasmid served as the template for PCR amplification of the apramycin resistance cassette, flanked by FRT sites and a 50bp overlap region with the target gene. Electrocompetent *K. pneumoniae* was transformed with pACBSR-hyg for Recombineering. The PCR amplified cassette was purified and electroporated into this strain with 1µg of DNA and plated on LB-apramycin. The pFLP-hygromycin plasmid was used to excise the antibiotic resistance cassette and all plasmids were cured from the strain through serial growth. PCR confirmed the mutant strains. See Key Resource Table and Supplemental Table 1 for plasmid information and primer sequences.

### Nitrogen and Carbon Sources Analysis

To determine sole carbon and nitrogen sources available to *K. pneumoniae*, the WT or Δ*ntrC* strains were growth overnight in M9 minimal media with ammonia and glucose, washed in PBS, diluted 1:1000 into M9 minimal media without ammonia (nitrogen testing) or without glucose (carbon testing). Overnight grown culture was diluted 1:1000 in sterile PBS and was aliquoted into Biolog Phenotype microarray 96-well plates PM1, PM2A (carbon testing), and PM3B (nitrogen testing). OD_600_ was determined via BioTek Epoch2 plate reader, grown for 24 hours shaking at 37°C. Area under the curve was calculated with software Prism v9.5.1 after subtracting background absorbance.

### Shotgun Sequencing and Analysis

DNA was extracted from stool using the Qiagen DNeasy PowerSoil Pro kit for the mouse experiments and Qiagen DNeasy PowerSoil for the cultivar experiments. Extracted DNA was quantified using the Quant-iTTM PicoGreen dsDNA assay kit (Thermo Fisher Scientific). Shotgun libraries were generated from 7.5 ng DNA using IDT for Illumina unique dual indexes and Illumina DNA Prep Library Prep kit for the mouse experiments and Illumina XT for the cultivar experiments at 1:4 scale reaction volume. Library success was assessed by Quant-iT PicoGreen dsDNA assay. Equal volumes of library were pooled from every sample and sequenced using a 300 cycle Nano kit on the Illumina MiSeq. Libraries were then repooled based on the demultiplexing statistics of the MiSeq Nano run. Final libraries were QCed on the Agilent BioAnalyzer to check the size distribution and absence of additional adaptor fragments. Libraries were sequenced on an Illumina Novaseq 6000 v1.5 flow cell for the mouse experiments and Illumina HiSeq 2500 v4 flow cell for the cultivar experiments, producing 2x150 bp paired-end reads. Extraction blanks and nucleic acid-free water were processed along with experimental samples to empirically assess environmental and reagent contamination. A laboratory-generated mock community consisting of DNA from Vibrio campbellii and Lambda phage was included as a positive sequencing control.

Shotgun metagenomic data were analyzed using Sunbeam version 2.1.1^52^. Quality control steps were performed by the default workflows in Sunbeam, which include removing adapters, reads of low sequence complexity and host-derived sequences. The abundance of bacteria was estimated using Kraken^53^. Reads were mapped to the KEGG database to estimate the abundance of bacterial gene orthologs^54^. Contigs were assembled from the samples in the FARMM dataset using MegaHit version 1.1.3^55^. The genes were predicted from the contigs using Prodigal version 2.6.3^56^. The predicted gene sequences were aligned against KEGG database using diamond aligner^57^.

Within sample diversity was assessed using Shannon diversity metric. Between sample similarity was assessed by Bray-Curtis distance. Community-level differences between sample groups were assessed using the PERMANOVA test. Differences in bacterial abundance or gene orthologs were assessed using linear models on log10 transformed relative abundances or RPKM values. Only bacteria with 1% mean relative abundance in at least one comparison are tested. P-values from multiple testing procedures will be corrected to control for a specified false discovery rate using Benjamini-Hochberg method.

### Metabolite quantification

Amino acids were quantified as previously described using a Waters Acquity uPLC System with an AccQ-Tag Ultra C18 1.7 μm 2.1x100mm column and a Photodiode Detector Array^34^. Fecal samples were homogenized in methanol (5 μL/mg stool) and centrifuged twice at 13,000g for 5 minutes. Amino acids in the supernatant were derivatized using the Waters AccQ-Tag Ultra Amino Acid Derivatization Kit (Waters Corporation, Milford, MA) and analyzed using the UPLC AAA H-Class Application Kit (Waters Corporation, Milford, MA) according to manufacturer’s instructions. All chemicals and reagents used were mass spectrometry grade.

Stool urea quantification was performed with the Quantichrom urea kit as follows: stool samples were homogenized in 10µL of ddH_2_O per mg stool. Solid debris was removed via centrifugation at 2500 x g for 10 minutes. Supernatant was tested for urea quantity per manufacturer protocol. A cutoff of 10% of the total microbiota, based on shotgun sequencing, was used for analysis of *K. pneumoniae* high and low categorization.

Metabolites were quantified by proton NMR as follows: Stool samples were resuspended and homogenized by vertexing. ^1^H NMR spectra were acquired on a Bruker Avance NEO 600-MHz spectrometer. Metabolites were assigned based on known spectral peaks and confirmed with a series of 2D NMR spectra.

### Quantification and Statistical Analysis

Prism Software (GraphPad, Inc.) was used to perform indicated statistical analyses. Statistical tests and p-values are noted in figure legends.

### Data and Code Availability

Shotgun sequencing data from the FARMM study is publicly available as previously reported in the Sequence Read Archive (SRA): PRJNA675301.

Shotgun sequencing data from this paper is publicly available in the SRA: PRJNA976029.

No original code was used for this study.

## Acknowledgements

This work was supported by the Division of Gastroenterology and Hepatology at the Hospital of the University of Pennsylvania and funded by the NIH P30 Center for Molecular Studies in Digestive and Liver Diseases, NIH R35GM139541, the Crohn’s and Colitis Foundation, and The Sherman Prize. We also acknowledge the assistance of the PennCHOP Microbiome Program, and the University of Pennsylvania Gnotobiotic Facility. We thank the Microbial Culture & Metabolomics Core of the PennCHOP Microbiome Program and the Center for Molecular Studies in Digestive and Liver Diseases (NIH P30DK050306) and the Penn Center for Nutritional Science & Medicine (PenNSAM) for their technical expertise. A.L.H. is supported by the Division of Gastroenterology basic science T32 (DK007066). We thank Lillian Chau, Lindsay Herman, and Dylan Curry for their technical expertise and assistance.

## Author Contributions

A.L.H, L.C.H., J.L., M.G., and G.D.W. designed, and analyzed the bacterial culture, human fecal testing, and mouse studies; A.L.H, L.C.H., J.L. performed the bacterial culture and mouse studies. E.S.F., J.F., L.L., and G.D.W designed the bioreactor studies; E.S.F and J.F. performed the bioreactor studies; C.T., V.T., and K.B performed the sequencing analysis. A.L.H, M.G., and G.D.W. wrote the manuscript.

---

**Supplemental Figure 1.**
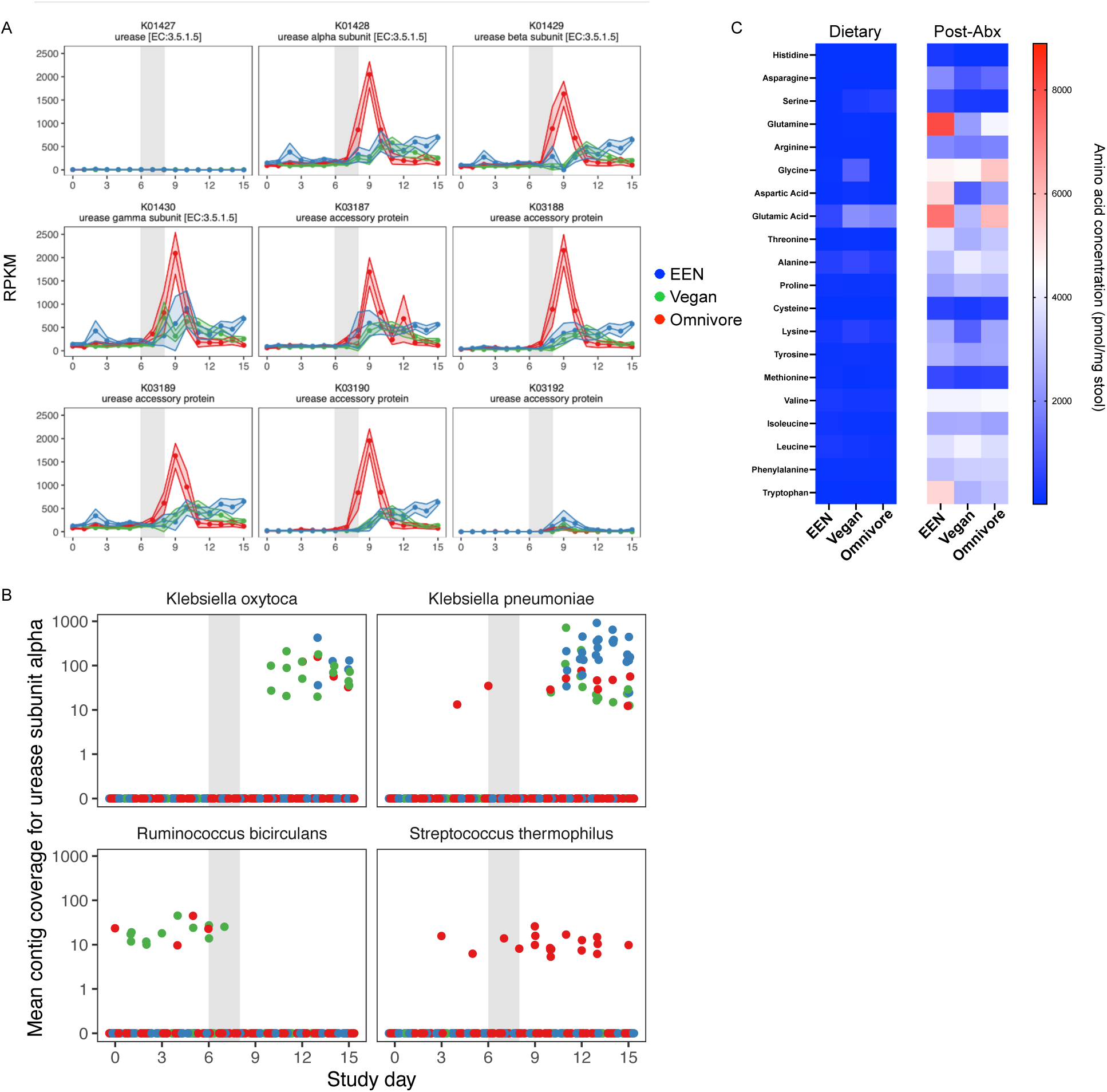
**A**. Urease gene abundance corrected for gene length and sequencing depth determined from shotgun metagenomic sequencing, organized by diet and study day. **B**. Urease operon reconstruction from shotgun metagenomic sequencing, with contig coverage displayed by study day and diet (EEN blue, omnivore red, vegan green). **C.** Amino acid quantification from the FARMM study on days 5 (dietary) and 9 (post-abx), as determined by HPLC, organized by dietary group. Data displayed as mean ± SE (**A**). n=10 subjects per dietary group.

**Supplemental Figure 2.**
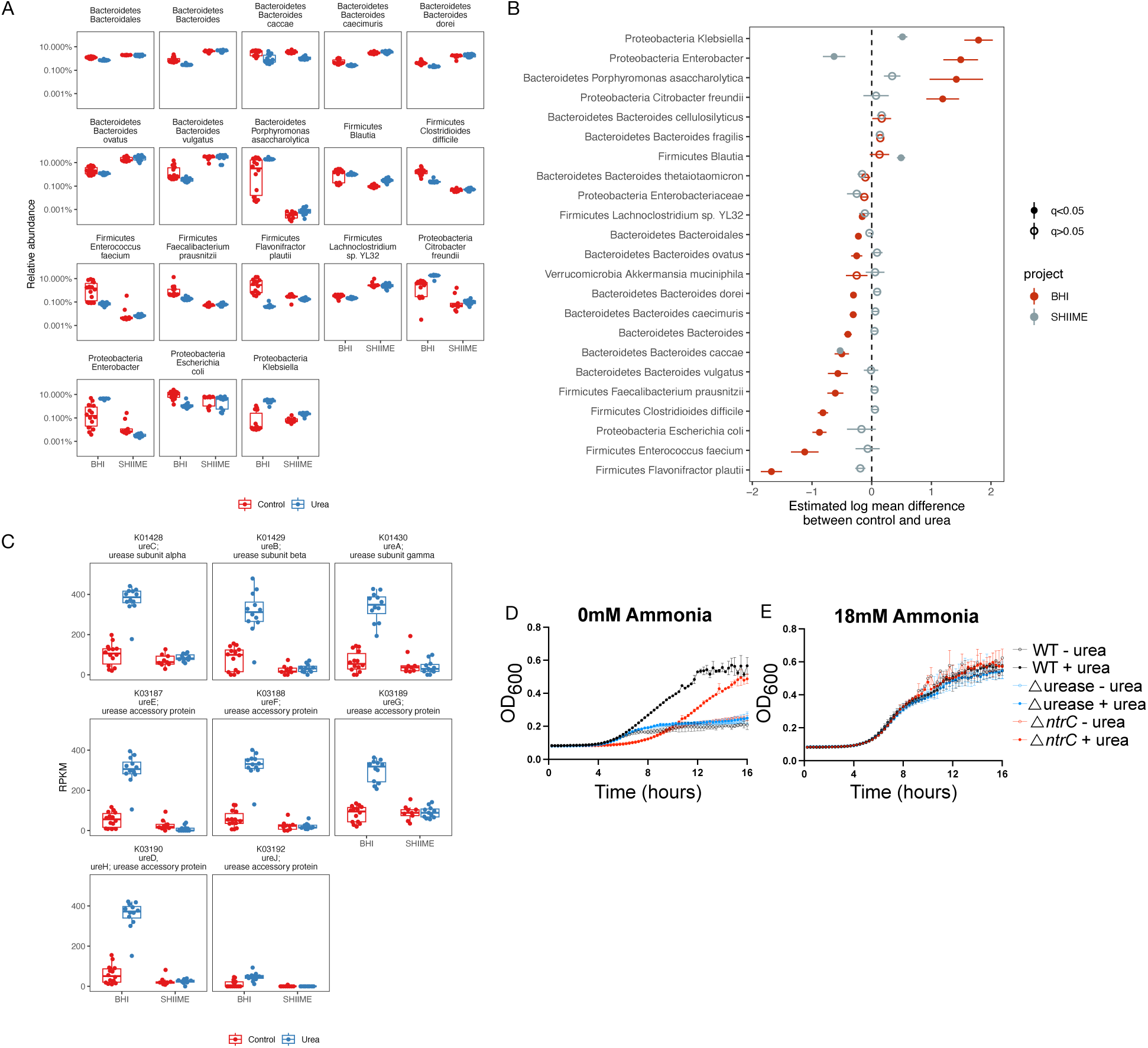
**A-C**. A human fecal sample was inoculated into a bioreactor system with BHI media or SHIME media with or without urea supplementation. Serial samples were collected and subjected to shotgun metagenomic sequencing revealing differences in species abundance (**A-B**) and urease gene abundance (**C**). **D and E**. WT *K. pneumoniae* and isogenic mutants of the urease operon and *ntrC* gene grown in M9 minimal media with 0mM (**D**), 18mM (**E**) ammonia with or without 5mM urea supplementation. OD_600_ was monitored during aerobic growth for 16 hours. Data presented as mean with quartiles noted by box and whisker plot (**A and C**), median and first and third quartiles (**B**), or mean ± SD (**D and E**); n= 15 time points per condition (**A-C**), or n= 3 wells per condition (**D-E**). Data representative of three independent experiments (**D-E**). Results of linear model on log_10_ transformed abundance.

**Supplemental Figure 3.**
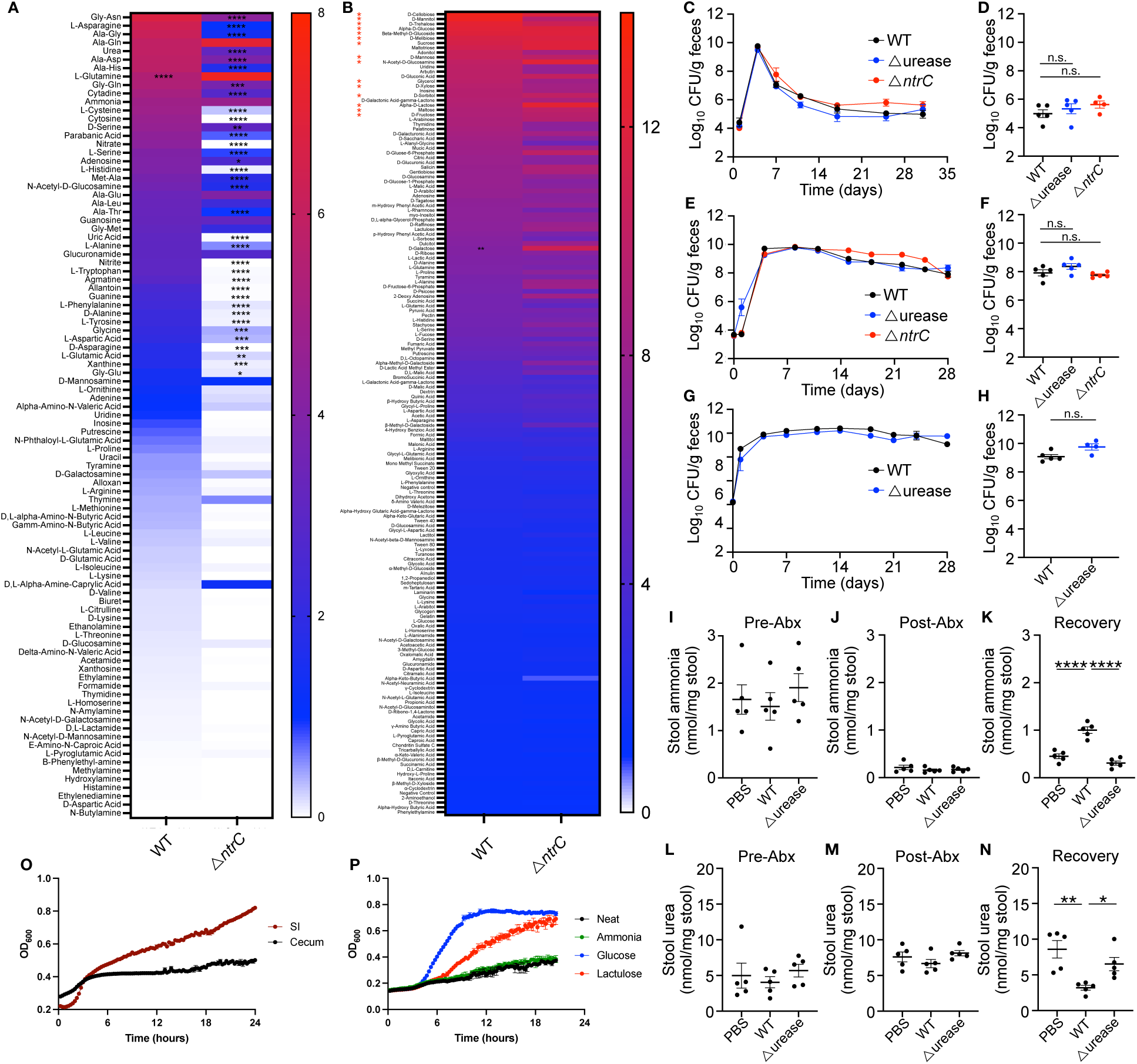
**A and B**. WT and Δ*ntrC K. pneumoniae* were grown in M9 minimal media with single nitrogen sources (**A**) or carbon sources (**B**) as indicated. Heat map represents area under the curve of OD_600_ monitored during aerobic growth for 24 hours. Red asterisks denote simple or alcohol sugars in the top 25 listed compounds. **C-H**. Mice were provided a standard chow diet (**C and D**), a low protein diet (**E and F**), or a fiber free diet (**G and H**) and gavaged with WT, Δ*ntrC,* or Δurease *K. pneumoniae* after abx pre-treatment and fecal CFU was monitored for the subsequent 4 weeks. (**C, E, and G**). CFU quantification 4 weeks after gavage for each respective diet (**D, F, and H**). **I-K**. Fecal ammonia was quantified in mice colonized with WT, Δ*ntrC,* or Δurease *K. pneumoniae* before abx (**I**), after abx (**J**) and 3 days after gavage (**K**). **L-N**. Fecal urea was quantified in mice colonized with WT, Δ*ntrC,* or Δurease *K. pneumoniae* before abx (**L**), after abx (**M**) and 3 days after gavage (**N**). **O and P**. *Ex-vivo* growth of *K. pneumoniae* in SI or cecal material (**O**) or in cecal material supplemented with ammonia, glucose, or lactulose (**P**) under anaerobic conditions. Data presented mean ± SEM (**C-P**), n=5 mice per group (**C-N**), or n=3 wells per experiment (**A, B, O, P**). Data represents two to three independent experiments. Results of one-way ANOVA with Bonferroni correction for multiple comparisons, n.s., not significant, n.s., not significant, *p<0.05, **p<0.01, ***p<0.001, ****p<0.0001.

**Supplemental Figure 4.**
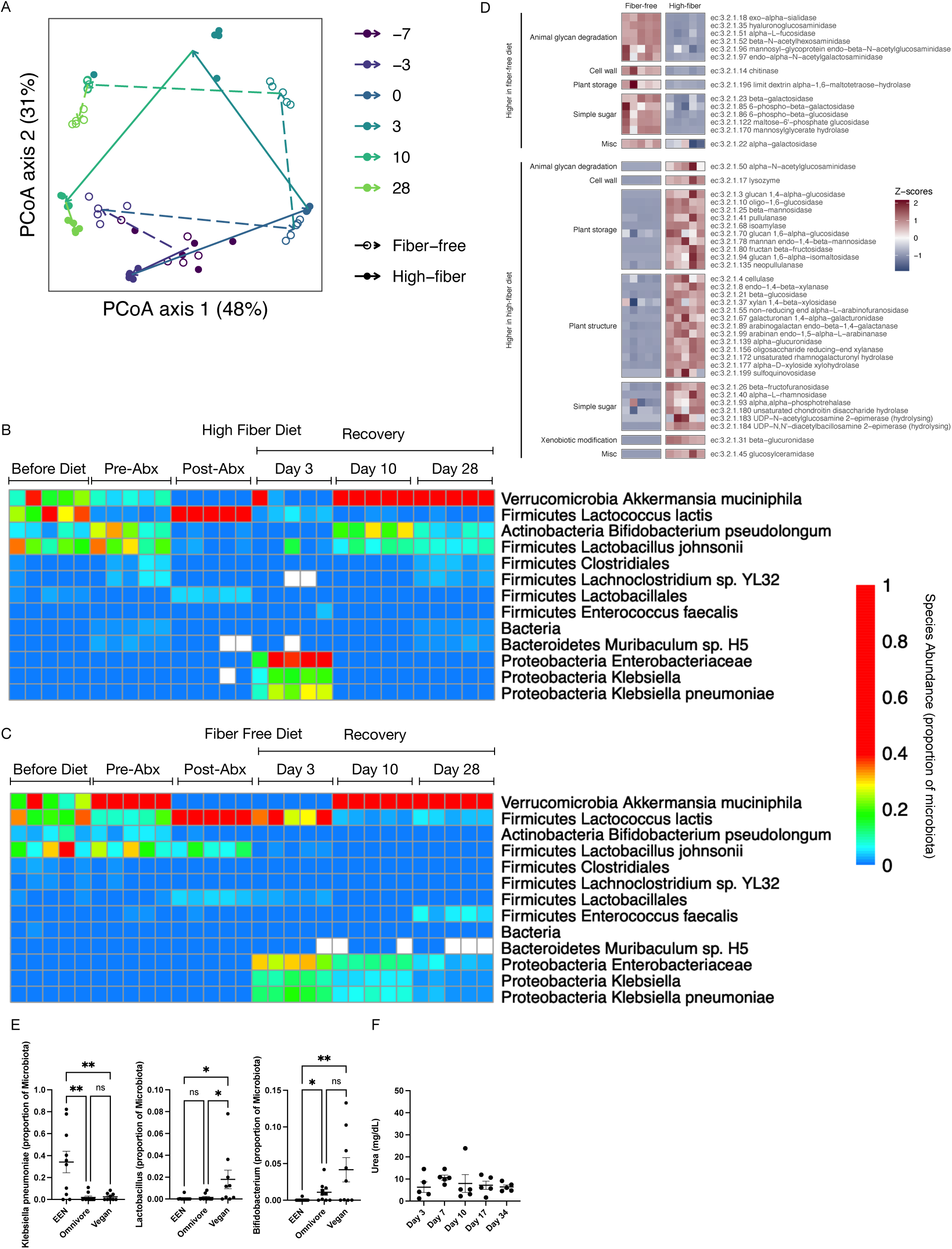
**A-D.** Mice were provided a HF or FF diet, treated with antibiotics, and gavaged with *K. pneumoniae*. PCoA plot of Bray-Curtis distances from mouse shotgun metagenomic sequencing from HF and FF dietary groups over time, as denoted by color. Each mouse is indicated by a single point with lines and arrows connecting the centroids of consecutive timepoints of the study (**A**). Heat map of species abundance (**B and C**) and Z-scores of glycoside hydrolase abundance (**D**) as determined by shotgun metagenomic sequencing over the course of the study period, with each mouse represented by a single column. **E.** Species abundance of *K. pneumoniae, Lactobacillus*, and *Bifidobacterium* from the FARMM trial shotgun metagenomic sequencing, organized by dietary group. **F.** Urea quantification from mice on a HF diet after *K. pneumoniae* gavage. Data presented as mean ± SD (**E and F**), n=5 mice per group (**A-D and F**), n=10 subjects per dietary group (**E**). Results of one-way ANOVA with Bonferroni correction, n.s. not significant, *p<0.05, **p<0.01 (**E and F**).

